# An Ultrasensitive Luciferase Reporter Reveals Low-Level Extrahepatic mRNA Expression for Enhanced Apparent Biodistribution of Lipid Nanoparticles

**DOI:** 10.64898/2026.09.15.751785

**Authors:** Vince Cataldi, Samrat Chakraborty, Adam W.G Alani

## Abstract

Reporter mRNAs are widely used to screen lipid nanoparticle (LNP) formulations and characterize functional mRNA expression in vivo, yet conventional firefly luciferase (FLuc) imaging fails to detect low-level expression. Consequently, weak or absent FLuc signal can be misinterpreted as failed LNP delivery or translation, potentially excluding formulations and overlooking extrahepatic or off-target expression. Here, we compared FLuc/D-luciferin with an ultrasensitive NLuc–GFP bioluminescence resonance energy transfer luciferase reporter detected with fluorofurimazine (FFz), using SM-102 and U105 LNPs following intravenous administration in mice. Both reporters showed predominantly hepatic and splenic expression. However, NLuc–GFP/FFz produced substantially greater photon output and detected reporter activity in the brain, heart, kidney, lymph nodes and other tissue where FLuc activity was weak or undetectable. Collectively, we show traditional FLuc imaging can underappreciate mRNA delivery, not necessarily because LNPs fail to reach or deliver mRNA to the tissue but because low-level FLuc expression may fall below its assay sensitivity. This limitation can underappreciate mRNA delivery and unintended protein expression in non-target organs. Therefore, we describe a reproducible workflow encompassing reporter design, LNP formulation and characterization, controlled reporter comparison, ex vivo organ imaging, and a cost-effective DSPE–PEG2000-micelle FFz formulation. This provides a more sensitive framework that improves detection and interpretation of low-level functional expression during preclinical mRNA-LNP screening and safety assessment.

## 1. Introduction

The investigation of nanoparticles for mRNA delivery requires reliable, sensitive techniques to evaluate the in vivo apparent biodistribution and subsequent protein expression. Although progress in lipid nanoparticles (LNPs) and mRNA chemistry has greatly enhanced delivery efficiency, interpreting results from LNPs and other mRNA delivery systems heavily depends on reporter constructs, such as luciferases or fluorescent probes, to assess tissue delivery and protein expression. In some studies, barcoding approaches have been introduced to increase screening throughput as well as infer LNP biodistribution; however, these methods remain relatively limited, and reporter mRNAs are still commonly used alongside, as barcodes cannot predict functional mRNA expression and so comparing top-performing delivery candidates often relies on luciferase reporters (1–5). Therefore, this work establishes luciferase reporter design and selection to be considered experimental variables in mRNA delivery. Specifically, a poorly optimized reporter can underestimate apparent delivery performance and obscure whether the absence of signal is due to inefficient delivery, poor mRNA translation, insufficient reporter sensitivity, or even substrate bioavailability (6–10).

Firefly luciferase (FLuc), first cloned in the late 1980s, rapidly became a gold-standard reporter for gene expression because of its novelty, ease of use, and compatibility with in vitro and in vivo imaging applications (11, 12). Despite its widespread use, FLuc reporter systems have several limitations for illuminating information for mRNA delivery studies. FLuc produces fewer photons than newer engineered luciferase systems and requires ATP, oxygen, magnesium, and exogenously administered D-luciferin, making bioluminescent output dependent on both mRNA expression and tissue substrate bioavailability (11, 12). Radiotracer pharmacokinetic studies have demonstrated low cellular uptake and rapid efflux of D-luciferin, with relatively low substrate accumulation in skeletal muscle, myocardium, and brain; overall uptake in the liver, lungs, myocardium, and skeletal muscle being much lower following intraperitoneal administration than intravenous administration (13). Another study using ^14^C-labeled D-luciferin confirmed heterogeneous tissue distribution, early renal elimination, and low brain uptake, with intraperitoneal administration producing slower and lower peak blood and organ concentrations than intravenous administration (14). Although intraperitoneal administration remains widely used because of its simplicity, direct comparison demonstrated that intravenous D-luciferin produced 5.6-fold higher peak photon emission, a shorter and less variable time to peak signal, and better measurement repeatability, yet reported literature still heavily relies on intraperitoneal (13, 14) comparisons. Consequently, conventional FLuc reporting can underestimate low-level mRNA delivery and subsequent protein expression, with insufficient detection causing promising formulations to be discarded and extrahepatic or off-target protein expression to be overlooked particularly in tissues with limited D-luciferin availability, limiting reporter expression across animals and organs which can misinterpret the apparent biodistribution of LNP-mRNA delivery

More recently, NanoLuc luciferase (NLuc) has emerged as a powerful alternative reporter for mRNA delivery studies because of its high intrinsic brightness, ATP-independent activity, and efficient substrate chemistry compared with conventional FLuc systems. (15). Ferraresso et al. employed NLuc mRNA in a swine model to show that a known ionizable LNP formulation (ALC-0315) generated detectable protein expression across multiple major organs after intravenous injection. (16). These findings highlight that reporter sensitivity can significantly affect the interpretation of mRNA biodistribution and imply that extrahepatic translation might be more prevalent than typically detected with FLuc. However, that study only assessed a single LNP formulation in a large-animal model and did not include a direct comparison with FLuc mRNA. Since most initial formulation screenings are conducted in mice, where low off-target translation may be near the detection threshold of conventional FLuc reporters or substrate availability is uncertain, systematic comparisons utilizing more sensitive mRNA reporters are essential to assess whether mRNA expression varies among LNP systems and to ensure that biodistribution is not consistently underestimated. An important advancement toward improving reporter sensitivity was reported by Schaub et al., who developed a bright and sensitive deep-tissue luciferase for in vivo imaging. The construct was a bifunctional reporter consisting of enhanced green fluorescent protein (eGFP) fused to NLuc (*nicknamed NLuc-GFP here)* (17). Oxidation of the NLuc substrate generates intramolecular *bioluminescence resonance energy transfer* (BRET), through which excitation energy is transferred from NLuc to GFP. This produces a green-shifted luminescent signal and increases detectable luminescence output while retaining GFP fluorescence, serving as a dual-purpose reporter for mRNA exploration. In Schaub et. al.’s work, the resulting luciferase reporter enabled highly sensitive longitudinal imaging in mice for deep-tissue imaging tumor cells in lymph nodes and embedded tissue, ultimately providing improved signal performance relative to conventional or engineered luciferase reporters (17). In parallel with these improvements to NLuc reporters, Su et al. addressed limitations associated with the conventional NLuc substrate furimazine, whose poor aqueous solubility and bioavailability restrict its performance in vivo. Their development of fluorofurimazine (FFz), an in vivo-optimized furimazine analogue with improved solubility and bioavailability, substantially increased the brightness of NLuc-based reporters in mice (18, 19). Together, these advances established a complementary strategy in which NLuc–GFP improves reporter sensitivity and provides dual bioluminescent and fluorescent readouts. At the same time, FFz increases effective substrate delivery and detectable photon output.

Ultimately, the goal of this study was to demonstrate low-level mRNA expression by employing NLuc–GFP, along with an alternative micelle formulation of FFz, as an ultrasensitive reporter system to compare previously reported ionizable LNP formulations for mRNA/protein translation and biodistribution in the most commonly used small-animal model, *Mus musculus*. By applying a luciferase designed to provide greater sensitivity than FLuc, we sought to determine whether the magnitude and apparent biodistribution of mRNA translation differ among LNPs, including the clinically validated SM-102 ionizable lipid nanoparticle and a previously reported U105 lipid. The selection of the reporter emphasizes whether tissue expression differences have been underestimated when using the less sensitive FLuc reporter and its substrate. Additionally, since FFz is typically available as an expensive commercial product, we developed an in-house FFz formulation utilizing DSPE–PEG2000 micelles, which enabled high substrate encapsulation and could be used directly after hydration as a cost-effective alternative. Lastly, NLuc-GFP was engineered with a degron to enable transient and longitudinal bioluminescence imaging studies. This strategy combines reporter engineering with substrate formulation optimization through an accessible FFz delivery system, enhancing the sensitivity and practicality of mRNA delivery screening in the most common animal model.

## 2. Results

### 2.1 Synthesis and Characterization of NLuc-GFP

The initial phase focused on the design, synthesis, and characterization of an in vitro-transcribed NLuc-GFP reporter mRNA for an ultrasensitive in vivo screening tool. This construct was based on the BRET NLuc reporter originally described by Schaub et al. and was selected because of its enhanced signal output in deep tissue, improved suitability for in vivo optical imaging, and its retention of the native eGFP, which acts as a multi-purpose reporter mRNA (17). Sequence information corresponding to the NLuc-GFP fusion was retrieved from the NCBI nucleotide database and verified by BLAST alignment during design. The resulting construct was assembled in an IVT-compatible plasmid backbone containing a T7 promoter, CleanCap-compatible initiation sequence, optimized 5′ and 3′ untranslated regions like BioNTech’s 3’UTR, and a segmented poly(A) tail design (30A+30A+43A); see Supplemental Figure 1 (7, 20–24). As shown in Figure 1A, the DNA map shows the key elements required for mRNA synthesis. Following plasmid transformation, linearization, and sequence verification, NLuc-GFP mRNA was generated under co-transcriptional capping conditions and purified for downstream use. Initial quality assessment by NanoDrop confirmed successful RNA production with a high-purity absorbance profile and an A260/A280 ratio of 2.04, consistent with relatively pure mRNA preparations (Figure 1B). To further assess transcript integrity and size distribution, the sample was analyzed by capillary electrophoresis using a Bioanalyzer. The resulting virtual gel image showed a single discrete product at the expected size (∼1900 bp), while the electropherogram displayed one dominant peak with minimal evidence of fragmentation or lower-molecular-weight byproducts (Figure 1C). Together, these data confirmed that the NLuc-GFP construct could be reproducibly synthesized as a clean, high-purity IVT mRNA suitable for the subsequent sensitive in vivo screening applications.

**Figure 1:**
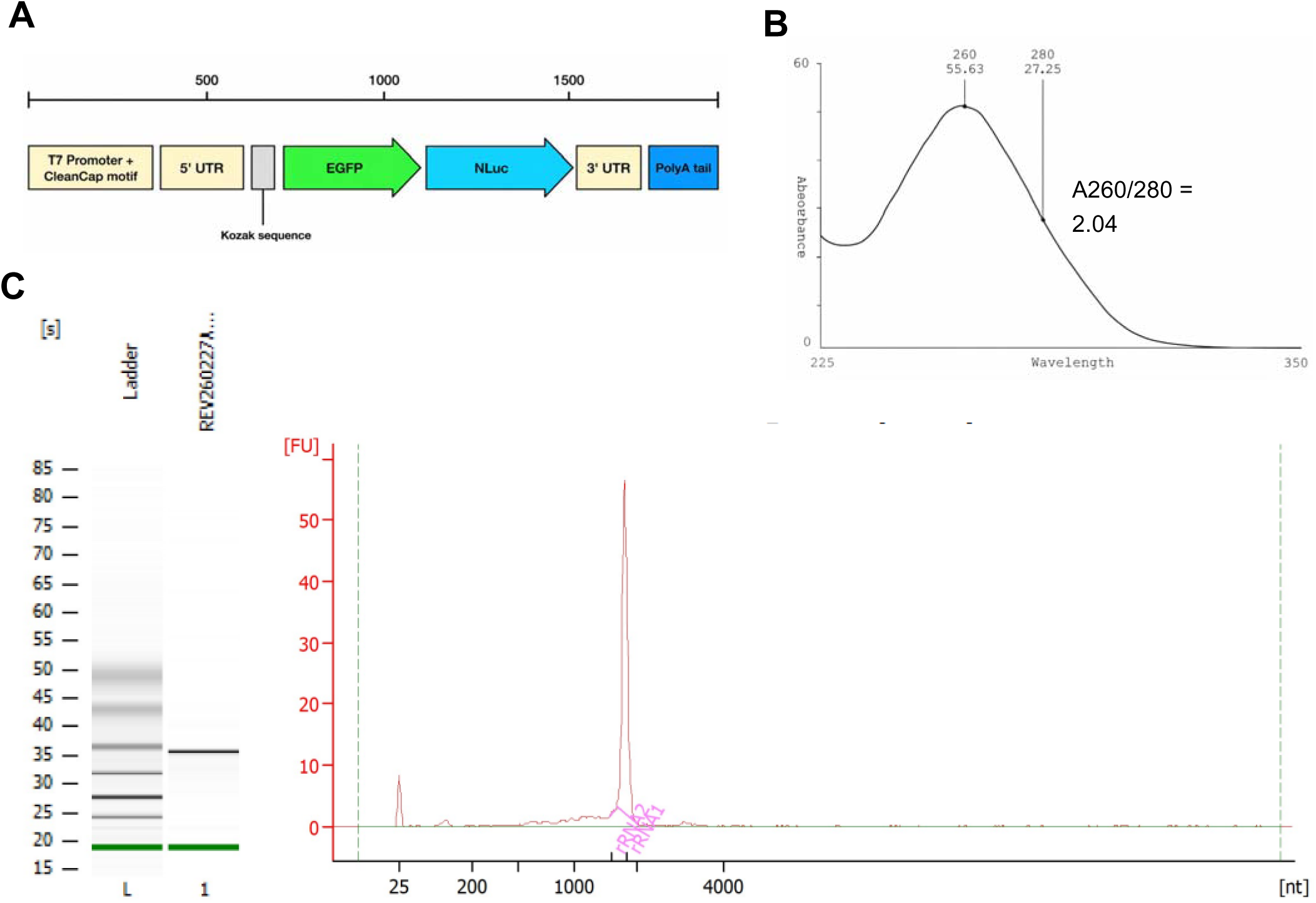
Design and Characterization of the NanoLuc–GFP mRNA construct. (A) Schematic of the in vitro–transcribed NLuc–GFP mRNA construct highlighting elements, including the 5′ untranslated region (UTR), coding sequence, 3′ UTR, and poly(A) tail. (B) NanoDrop absorbance spectrum of purified mRNA demonstrating high purity, with an A260/280 ratio > 2.0, indicative of minimal protein contamination. (C) Bioanalyzer assessment of mRNA integrity: capillary electrophoresis (left) shows a single discrete band corresponding to the NLuc–GFP transcript alongside a molecular weight ladder, while the electropherogram (right) displays a single sharp peak, confirming a homogeneous and intact mRNA population.

### 2.2 Formulation of DSPE-PEG2k Fluorofurimazine

To improve the delivery of fluorofurimazine (FFz) and reduce reliance on commercially available substrate formulations or on variability from ex vivo-based solutions. FFz was formulated into inexpensive DSPE– PEG2000 micelles using thin-film hydration. A DSPE–PEG2000:FFz mass ratio of 10:1 produced a homogeneous dispersion following hydration in 5% dextrose. Dynamic light scattering demonstrated that the FFz-loaded micelles formed a predominantly nanoscale population with a mean hydrodynamic diameter of 243 nm and a polydispersity index of 0.23 (Supplemental Figure 2A). The formulation exhibited a zeta potential of –23 mV (Supplemental Figure 2B). Unfortunately, once FFz is in an aqueous solution, it slowly loses stability, with around a <15% decrease in FFz concentration in 8 hours at room temperature. To extend the formulation’s shelf life, the micelles were lyophilized; upon reconstitution, the lyophilized powder yielded micelles with similar FFz loading and comparable size and polydispersity (Supplemental S2A-B). These data indicate that the formulation maintained both colloidal and chemical stability throughout the evaluated period.

Quantification of disrupted micelle samples showed approximately 95% encapsulation efficiency and recovery of FFz, with the remainder as free, soluble FFz, indicating minimal substrate loss during film formation, hydration, and sample processing. However, centrifugal filtration or post-processing of micelles is unnecessary, and the FFz-micelles can be used directly after hydration, as all FFz remains in solution. Collectively, these results demonstrate that DSPE–PEG2000 micellization enabled efficient aqueous dispersion of FFz while producing a relatively uniform and stable formulation that can be freeze-dried

### 2.3 Formulation and characterization of U105 and SM-102 lipid nanoparticles

To investigate ionizable lipid differences in apparent mRNA biodistribution, LNPs were prepared using either SM-102, a clinically validated ionizable lipid used in approved mRNA vaccine formulations, or U105, a non-clinical ionizable lipid previously reported by Liu et al.(25). Each lipid was formulated separately with FLuc (TriLink) or NLuc-GFP mRNA, producing four formulations: SM-102-FLuc, SM-102-NLuc-GFP, U105-FLuc, and U105-NLuc-GFP. All formulations used the same 50:35:12.5:2.5 molar ratio of ionizable lipid:cholesterol:DOPE:DMG-PEG2000 (Figure 2A). The structures of SM-102 and U105 are shown in Figure 2B. LNPs were made by microfluidic mixing using the classic workflow (Figure 2C). All four formulations produced nanoscale particles with similar physicochemical properties (Figure 2D). SM-102-FLuc and SM-102-NLuc-GFP measured 95 and 91 nm, with PDIs of 0.15 and 0.18, respectively. U105-FLuc and U105-NLuc-GFP measured 78 and 84 nm, with PDIs of 0.17 and 0.21. Zeta potential was slightly negative for all formulations, ranging from −5.2 to −4.1 mV. Encapsulation efficiencies, as measured by Ribogreen assay, indicated encapsulation exceeding 90% for all formulations tested. These results confirm that both reporter mRNAs were efficiently encapsulated within SM-102 and U105 LNPs. The comparable size, PDI, surface charge, and encapsulation efficiency of the four formulations supported their subsequent use for comparing reporter-dependent detection and ionizable lipid-dependent mRNA biodistribution in vivo.

**Figure 2.**
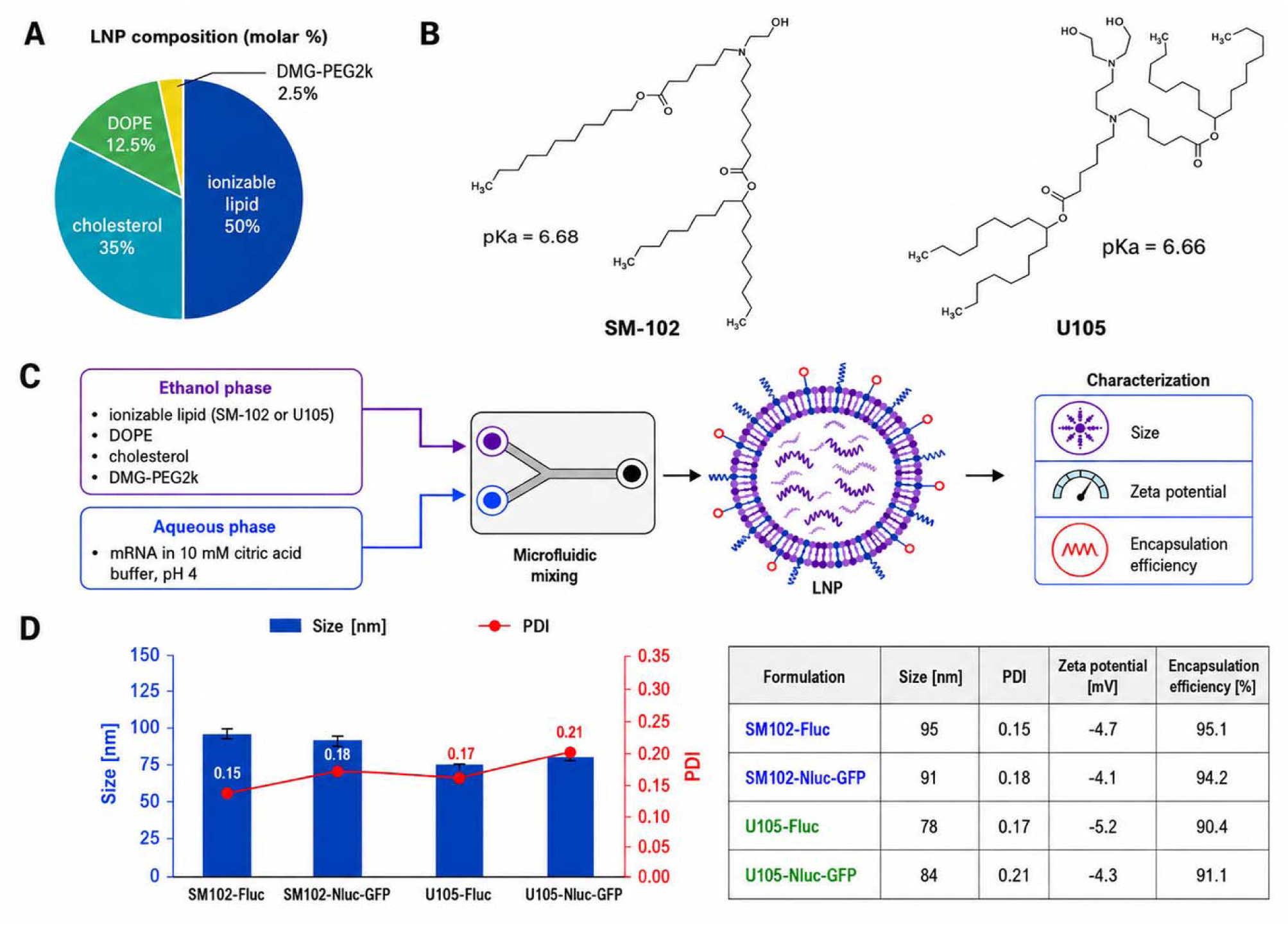
Formulation and characterization of SM-102 and U105 mRNA lipid nanoparticles. (A) LNP composition used for formulations. (B) Chemical structures of the two ionizable lipids evaluated: the clinically relevant SM-102 and the investigational lipid U105. (C) Schematic of the microfluidic formulation workflow. (D) Physicochemical characterization of the resulting LNPs by dynamic light scattering, zeta potential, and RiboGreen encapsulation assay.

### 2.4 NLuc-GFP reveals Low-Level mRNA Biodistribution Compared to FLuc

To assess how the luciferase reporter’s sensitivity affects the interpretation of nanoparticle uptake and apparent biodistribution of systemic mRNA/protein, LNPs containing mRNAs encoding either NLuc-GFP or codon-optimized FLuc (TriLink) were administered intravenously at 5 µg of mRNA per animal. Whole-body bioluminescence imaging and ex vivo organ biodistribution were performed 6 h post-injection to capture early/maximum reporter expression. Because FLuc and NLuc-GFP require different substrates, signal intensity was interpreted as a measure of reporter sensitivity under the typical imaging conditions recommended by the supporting literature (2, 16, 17, 24–27), rather than as a direct substrate comparison of the translated protein and assuming that each amount of mRNA produces similar protein respectively. Together the NLuc-GFP/FFz system is dramatically more sensitive, for reference NLuc alone is 100-fold more sensitive than FLuc, but with NLuc coupled with the resonance transfer from eGFP, it produces luminescence that penetrates tissue better than that of classical NLuc and outperforms engineered firefly luciferase or NLuc.. Due to the sensitivity involved, the substrates FFz and D-luciferin (D-luc) are administered at different doses. The FFz injection was standardized at 200 µg per mouse (0.468 µmol), whereas 3 mg of D-luciferin (9.38 µmol) was used, representing a twentyfold difference in substrate amounts. This clearly demonstrates the sensitivity in a manner we consider appropriate for comparison and as seen in the literature (2, 26–30)

The first formulation evaluated was the U105 LNP, a commercially available, non-clinically approved ionizable lipid formulation used as an investigational LNP platform (Figure 3). U105 was selected because prior work by Liu et al (25) demonstrated classic hepatic mRNA whole-body expression following systemic delivery, but detailed ex vivo organ biodistribution was not extensively characterized. Therefore, U105 provided an opportunity to evaluate further how this non-clinical ionizable lipid drives reporter expression across major organs and to determine whether NLuc-GFP could reveal tissue expression that may be underestimated with a conventional FLuc reporter. We detected whole-body bioluminescence signals after 6h detected from both luciferase mRNAs, with NLuc-GFP producing a substantially stronger signal throughout the animal (Figure 3B–C), resulting in a 4.46-fold difference in whole-body luminescence, with the suspected hepatic tropism. This difference in reporters was evident during ex vivo organ imaging, where NLuc-GFP showed signal in all harvested organs, including the liver, lungs, spleen, kidney, heart, and brain (Figure 3D). In contrast, the FLuc signal showed lower luminescence intensity and appeared more restricted and near-background, especially in areas such as the heart, kidneys, and, mainly, the brain, where its substrate, D-luciferin, is often restricted. Quantitative analysis confirmed significantly greater NLuc–GFP luminescence in every organ examined, ranging from 20– to 112-fold above FLuc (Figure 3E). The largest differences occurred in the brain (112-fold), lungs (67-fold), and liver (50-fold), followed by the heart (32-fold), kidneys (29-fold), and spleen (20-fold). Importantly, mice receiving FFz without reporter mRNA exhibited no detectable whole-body or ex vivo organ luminescence, with all tissues remaining at background levels (Supplemental Figure S3). These findings confirm that the enhanced NLuc–GFP signals were reporter-dependent and demonstrate that reporter sensitivity substantially alters the apparent functional biodistribution of U105 LNPs.

**Figure 3.**
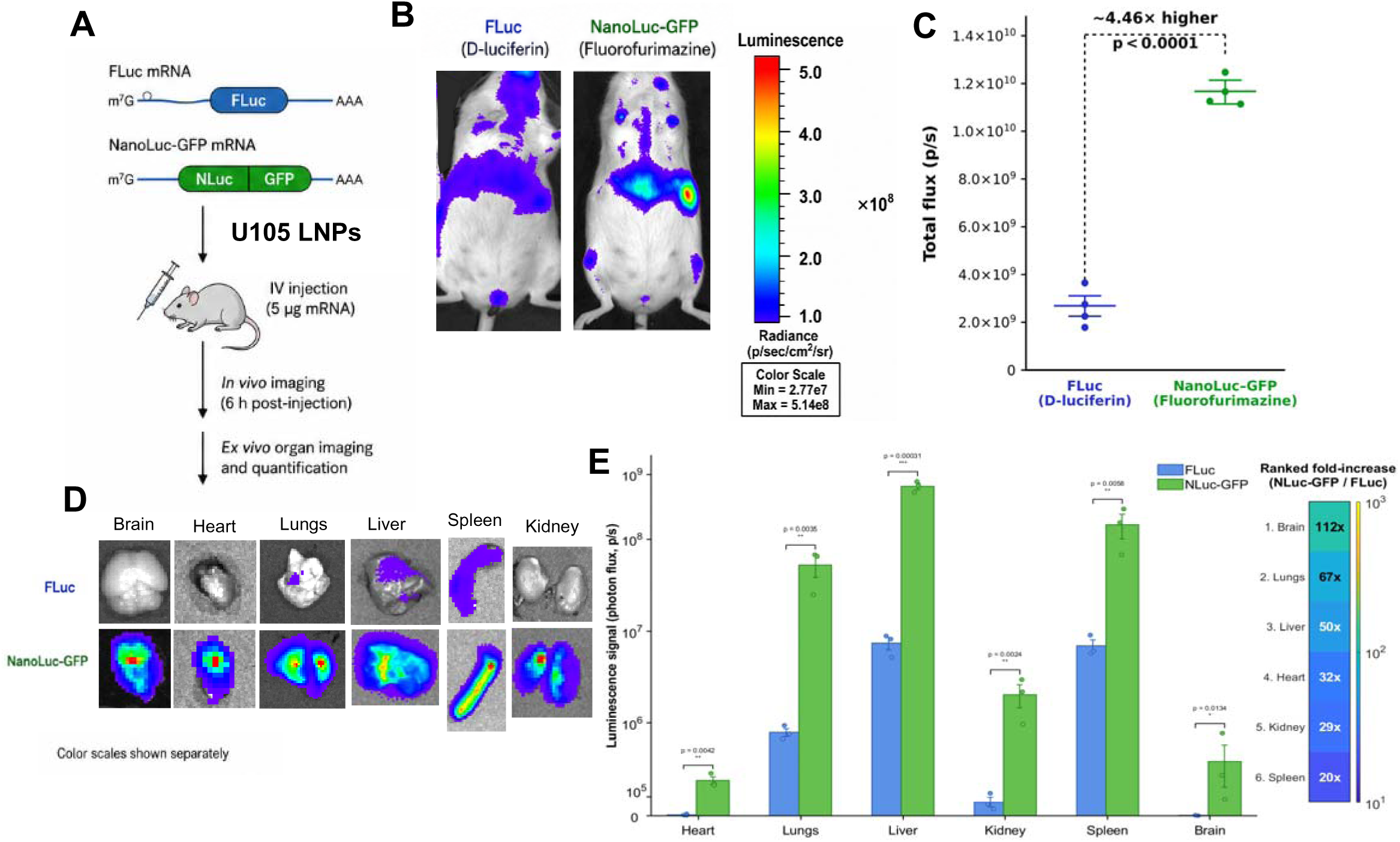
Bioluminescence following intravenous delivery of U105 LNPs. (A) Experimental design comparing U105 LNPs loaded with FLuc or NLuc-GFP mRNA, formulated under identical conditions and administered intravenously at 5 µg mRNA per mouse, followed by in vivo imaging and ex vivo organ analysis at 6 h post-injection. (B) Representative whole-body bioluminescence images showing stronger and broader signal for NLuc-GFP than FLuc following delivery by U105 LNPs. (C) Quantification of whole-body total flux demonstrating significantly higher signal from NLuc-GFP relative to FLuc. (D) Representative ex vivo organ bioluminescence images of brain, heart, lungs, liver, spleen, and kidney collected 6 h after injection. (E) Organ-level quantification showing that NLuc-GFP reveals greater signal across multiple tissues, including low-expression organs such as the brain, indicating that reporter sensitivity strongly influences the apparent biodistribution of U105 LNP-mediated mRNA delivery. Data is shown as mean ± SD (n=4)

To determine whether this reporter-dependent effect was specific to the U105 formulation or generalizable to other relevant LNP compositions, the same comparison was performed using the clinically approved SM-102 LNPs (Figure 4). SM-102 is a clinically validated ionizable lipid used in approved mRNA vaccine formulations, making it an important benchmark for evaluating whether reporter sensitivity influences biodistribution interpretation in a translational LNP platform. Similar to the U105 formulation, whole-body bioluminescence imaging showed a stronger detectable signal in animals receiving NLuc-GFP mRNA than in those receiving FLuc mRNA (Figure 4B–C). Ex vivo imaging again demonstrated that NLuc-GFP enabled clearer visualization of mRNA and protein reporter expression across multiple organs. In contrast, the FLuc signal remained comparatively weaker and less broadly resolved (Figure 4D), with tissue distribution limited to well-known tissues such as the liver and spleen, and faintly in the lungs. Quantification of ex vivo imaging further demonstrated luciferase reporter differences in apparent organ distribution. NLuc–GFP signal was detected throughout the brain, heart, lungs, liver, spleen, and kidneys, whereas FLuc signal was substantially weaker and appeared near background in the heart, kidneys, and brain (Figure 3D). This difference was particularly notable in the brain, where limited D-luciferin accessibility may further restrict FLuc detection. Quantitative analysis confirmed greater NLuc–GFP luminescence in every organ examined (Figure 3E). The fold difference was comparatively modest in the liver and spleen, at 2– and 3-fold, respectively, where dominant expression was readily detected using both reporters. In contrast, organs where expression is often weak, undetected, or interpreted as absent by conventional FLuc imaging showed dramatic increases with NLuc–GFP, including the kidneys (1,823-fold), brain (687-fold), and heart (536-fold), while the lungs showed a 32-fold increase. Importantly, mice receiving FFz without reporter mRNA exhibited no detectable whole-body or ex vivo organ luminescence, with all tissues remaining at background levels (Supplemental Figure S3). When FLuc was used, lower-expression tissues appeared weak or undetectable, potentially giving the impression that delivery was absent or highly restricted.

**Figure 4.**
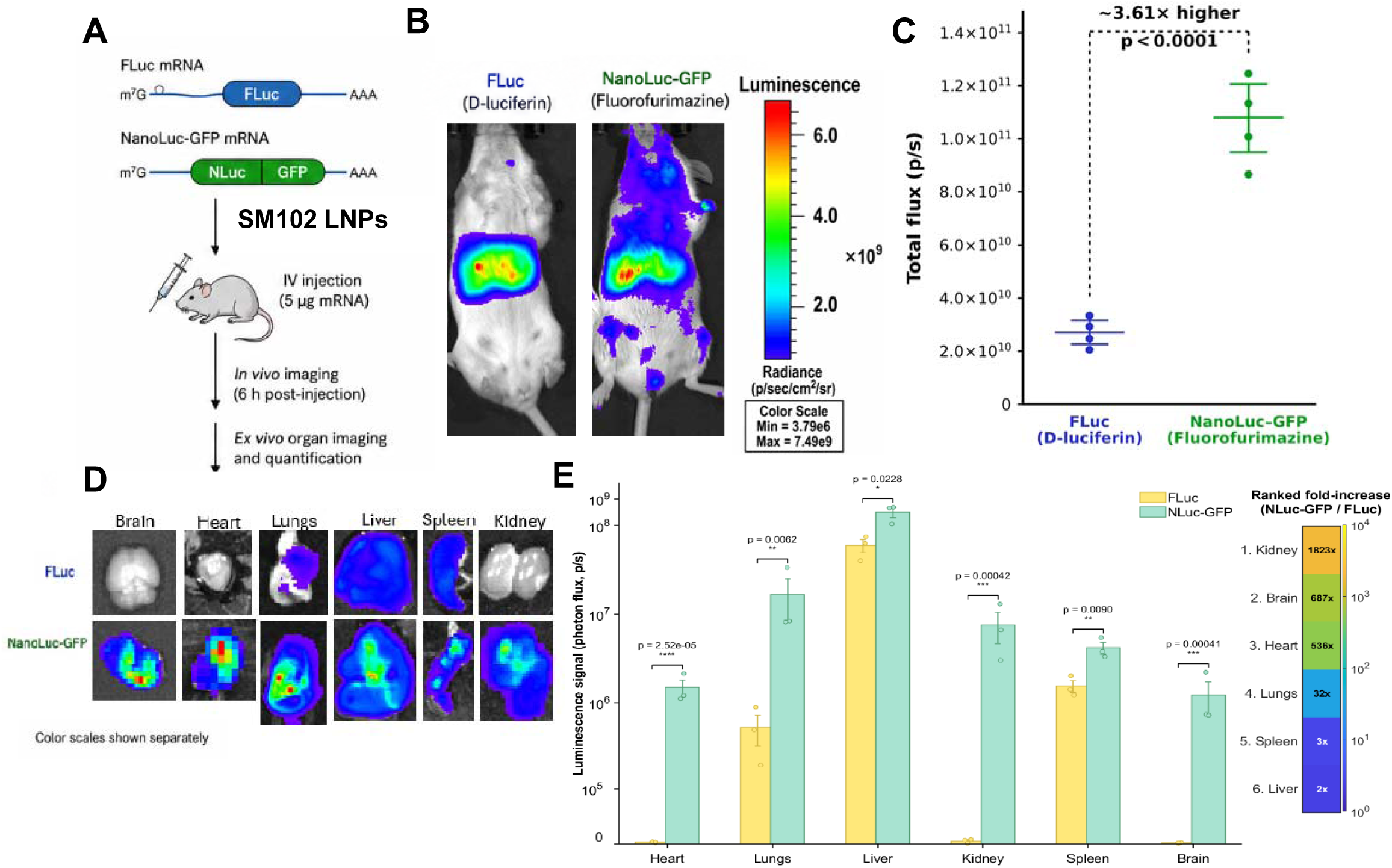
Bioluminescence following intravenous delivery of SM-102 LNPs. (A) Experimental design comparing SM-102 LNPs loaded with FLuc or NLuc-GFP mRNA, formulated under identical conditions and administered intravenously at 5 µg mRNA per mouse, followed by in vivo imaging and ex vivo organ analysis at 6 h post-injection. (B) Representative whole-body bioluminescence images showing stronger and broader signal for NLuc-GFP than FLuc following delivery by U105 LNPs. (C) Quantification of whole-body total flux demonstrating significantly higher signal from NLuc-GFP relative to FLuc. (D) Representative ex vivo organ bioluminescence images of brain, heart, lungs, liver, spleen, and kidney collected 6 h after injection. (E) Organ-level quantification showing that NLuc-GFP reveals greater signal across multiple tissues, including low-expression organs such as the brain, indicating that reporter sensitivity strongly influences the apparent biodistribution of U105 LNP-mediated mRNA delivery. Data is shown as mean ± SD (n=4)

In contrast, NLuc-GFP showed a measurable signal in tissues where FLuc approached background levels, indicating that mRNA delivery and translation were occurring but were below the detection threshold of the less sensitive FLuc reporter system. For both U105 and SM-102 LNPs, FLuc-associated brain signal was undetectable, whereas NLuc-GFP revealed measurable brain-associated signal, albeit low but still detectable, indicating nanoparticle uptake and mRNA translation. Therefore, this does not necessarily imply high brain delivery, but it demonstrates that low-level reporter expression can be missed when using the less sensitive FLuc reporter. Additionally, there was an indication that lymph nodes were being illuminated with the ultrasensitive luciferase. Therefore, apparent absence of FLuc signal should not be interpreted as definitive absence of nanoparticle/mRNA delivery or translation. Rather, in tissues with low expression, limited substrate access, or poor detectability, reporter sensitivity becomes the major determinant of whether expression can be detected at all. Together, the U105 LNP data in Figure 3 and the SM-102 LNP data in Figure 4 demonstrate that NLuc-GFP provides a more sensitive readout of LNP-mediated mRNA expression than FLuc across whole-body and ex vivo organ imaging. Additionally, there was no luminescence in the control mice injected with only FFz at the dose used in the mRNA experiments. These results support the use of NLuc-GFP as a higher-sensitivity reporter for evaluating mRNA delivery platforms, particularly when screening formulations or assessing low-signal tissues, where conventional FLuc imaging may underestimate the apparent biodistribution.

### 2.5 Long-Acting and Short-Acting Reporters

While the NLuc-GFP reporter substantially improved detection sensitivity and the apparent biodistribution of LNPs tested, its stability proved to be a notable variable. During early in vivo studies, NLuc-GFP expression persisted longer than expected (lasting> 7 days), and subsequent tests indicated that this prolonged signal was not due to continued mRNA translation but was most likely attributable to the long half-life (high stability) of the NLuc-GFP fusion protein after it had already been expressed. This property is beneficial for screening applications because it allows NLuc-GFP mRNA to be administered at almost any time and imaged reliably and flexibly over a broad window, with minimal signal loss. In this sense, the NLuc-GFP construct functioned as a “long-acting” reporter that was highly advantageous for maximizing sensitivity and enabling robust detection of delivery, particularly in tissues with relatively low expression. However, this same long half-life also introduced a limitation, because a persistent signal could obscure whether the expressed luminescence reflected newly synthesized protein or the retention of previously translated reporter. For studies aimed at comparing translation kinetics over time, a more transient reporter was needed. To address this limitation, a second NLuc-GFP construct was engineered to behave as a short-acting or destabilized reporter by incorporating a PEST degradation signal, as shown in Figure 5A. The long-acting and short-acting NLuc-GFP reporters are otherwise nearly identical in overall mRNA architecture (see Supplemental S1), with the critical difference being the addition of the PEST degron sequence to make a “short-acting construct.” PEST sequences are amino acid motifs enriched in proline (P), glutamic acid (E), serine (S), and threonine (T) that function as destabilizing elements within proteins. These motifs are recognized by cellular degradation machinery and promote rapid turnover, most commonly through proteasomal degradation (Figure 5A).

**Figure 5:**
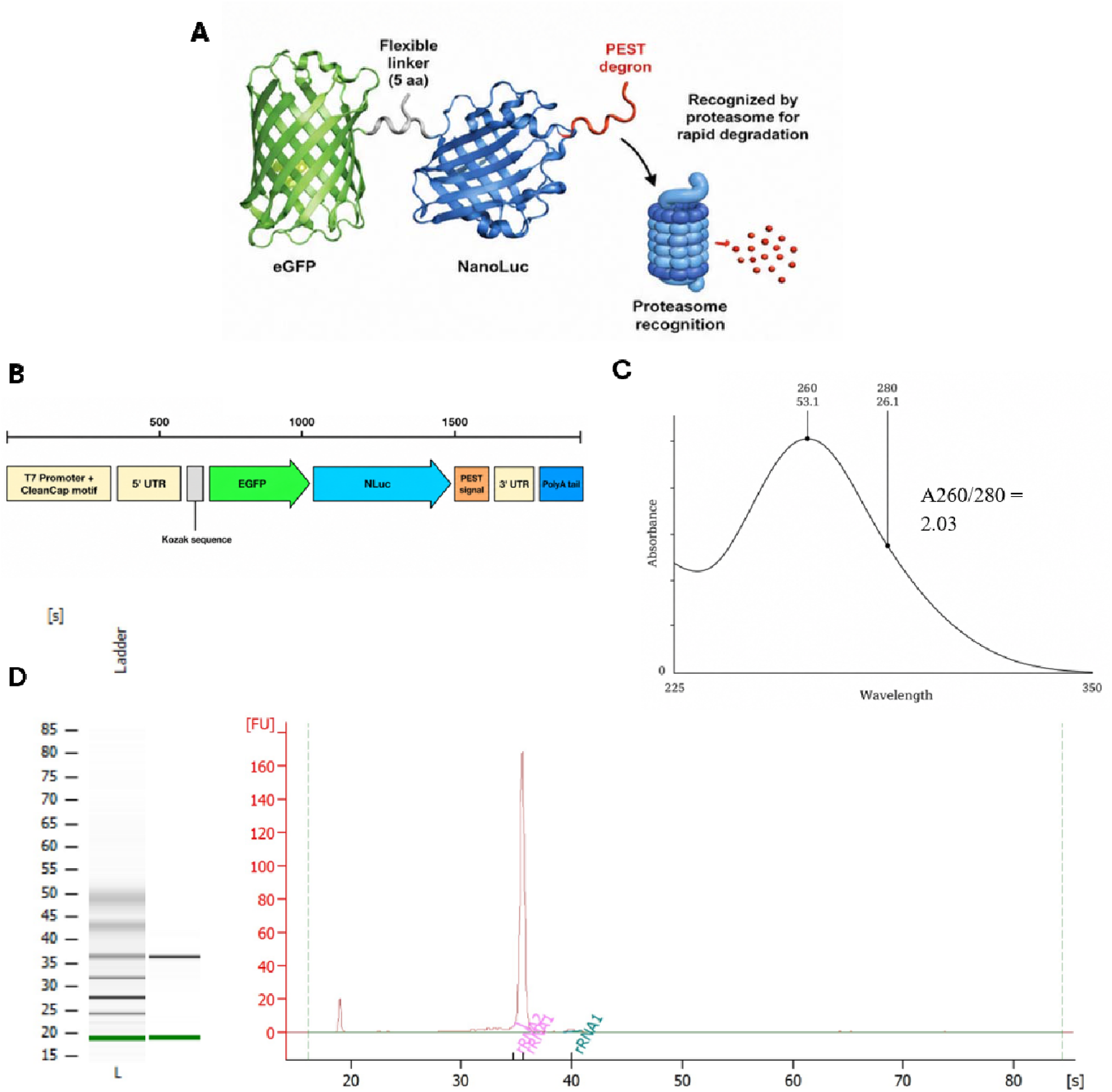
Design and Characterization of the NanoLuc–GFP-PEST mRNA construct. (A) Structural representation of the NLuc–GFP-PEST fusion protein. (B) Schematic of the in vitro–transcribed NLuc–GFP mRNA construct highlighting key elements. (C) NanoDrop absorbance spectrum of purified mRNA demonstrating high purity, with an A260/280 ratio > 2.0, indicative of minimal protein contamination. (D) Bioanalyzer assessment of mRNA integrity: capillary electrophoresis (left) shows a single discrete band corresponding to the NLuc–GFP transcript alongside a molecular weight ladder, while the electropherogram (right) displays a single sharp peak, confirming a homogeneous and intact mRNA population.

Incorporating the PEST sequence directs the NLuc-GFP fusion protein toward rapid intracellular degradation, lowering its half-life and diminishing signal persistence over time. This transformation shifts the reporter from a long-acting signal to a short-acting one, whose luminescence more accurately represents current translation rather than accumulated protein. The corresponding NLuc-GFP-PEST mRNA construct is shown in Figure 5B. Following in vitro transcription and purification, NanoDrop analysis showed an A260/A280 ratio of 2.03 (Figure 5C), and Bioanalyzer analysis demonstrated a single dominant band at the expected size (∼2010 bp) and an electropherogram peak with minimal evidence of artifacts (Figure 5D), confirming successful synthesis of a clean and intact NLuc-GFP-PEST mRNA for downstream in vivo use.

The functional consequences of PEST incorporation are evident as outlined in Figure 6; both NLuc-GFP and NLuc-GFP-PEST mRNAs were formulated under identical conditions with the clinically relevant SM-102 LNPs, administered by intravenous injection at 3 µg mRNA, and imaged longitudinally at 6 h, 24 h, 48 h, and 7 d post-injection Figure 6A. Whole-animal bioluminescence imaging (Figure 6B) showed that both constructs generated a strong early signal, confirming that the PEST-containing reporter remained fully capable of reporting successful translation shortly after delivery. However, unlike the long-acting NLuc-GFP reporter, the NLuc-GFP-PEST signal declined rapidly. Quantification of total luminescence (Figure 6C) confirmed this pattern, with the long-acting NLuc-GFP construct maintaining high signal across the measured time points, whereas the short-acting NLuc-GFP-PEST construct showed a marked decrease between 6, 24, and 48 h and reached background by 7 d. Together, these reporters provide a complementary reporter for evaluating mRNA delivery. Specifically, the long-acting NLuc-GFP construct offers maximum sensitivity and flexible screening capabilities. In contrast, the short-acting NLuc-GFP-PEST construct provides better resolution of translation kinetics, allowing examination of mRNA and protein production over time rather than at a single time point, while maintaining the apparent biodistribution in all organs (Supplemental S4). As a pair, they provide a more complementary reporter for understanding both the magnitude and duration of mRNA translation following delivery.

**Figure 6.**
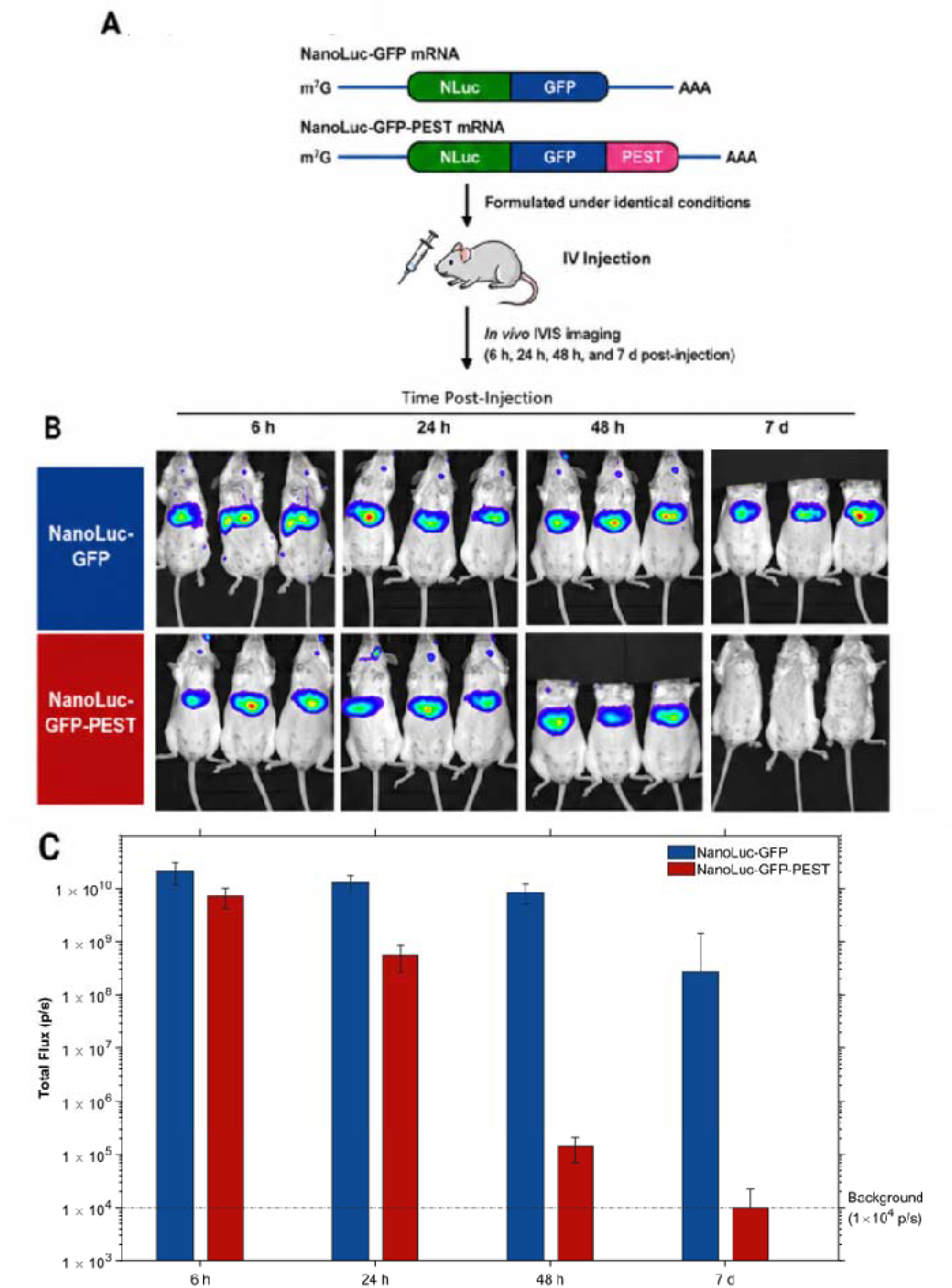
In vivo comparison of NanoLuc-GFP and NanoLuc-GFP-PEST reporter mRNA. (A) Experimental design comparing NanoLuc-GFP and destabilized NanoLuc-GFP-PEST mRNAs formulated under identical conditions and administered intravenously at 3 µg per mouse. In vivo IVIS imaging was performed at 6 h, 24 h, 48 h, and 7 d post-injection. (B) Representative whole-body bioluminescence images showing sustained NanoLuc-GFP signal compared with rapidly declining NanoLuc-GFP-PEST signal. (C) Quantification of total flux over time. NanoLuc-GFP maintained detectable expression through 7 d, whereas NanoLuc-GFP-PEST declined to background by 7 d. Data is shown as mean ± SD (n=3)

## 3. Discussion

FLuc remains widely used because it is well characterized and compatible with established imaging workflows. However, its output depends on reporter abundance and tissue-dependent D-luciferin bioavailability and distribution (9, 10). Radiotracer studies have demonstrated relatively low cellular uptake and rapid efflux of D-luciferin, along with heterogeneous tissue distribution with limited accumulation in the myocardium, skeletal muscle, bone, and brain (11, 12). Further, D-luciferin is also rapidly cleared through the kidneys, creating a time-dependent window of substrate availability. These limitations vary depending on the administration technique, with intraperitoneal administration being the most popular and showing significantly reduced biodistribution in tissue, with a 5.6-fold decrease in photon output. This is increasingly important when luciferase expression is low, because limited or transient substrate exposure may prevent the luciferase reporter from producing sufficient photon output to be detectable by IVIS. Weak FLuc signal may therefore reflect the combined effects of low luciferase expression and substrate pharmacokinetics rather than the complete absence of mRNA delivery or translation. Specifically, the brain, kidney, and heart represent the most prominent example of reporter-dependent detection in this study. FLuc activity in the brain was undetectable under the conditions evaluated, whereas NLuc-GFP produced measurable brain and other tissue signal with both U105 and SM-102 LNPs (Fig3 and 4). The limited uptake of D-luciferin can further restrict FLuc detection when reporter expression is already near the lower end of the measurable range (11, 12). Consequently, the absence of FLuc signal in the brain cannot independently establish that LNP delivery or mRNA translation did not occur. Expression may instead remain below the practical sensitivity of the FLuc/D-luciferin system and become detectable only with a brighter reporter and a more favorable substrate.

Respectfully, the brain-associated signal and other low-level expression should nevertheless be interpreted as an apparent biodistribution, as they do not independently establish specific transfection of select cell types. The signal may originate from vascular-associated expression, endothelial cells, or a certain population of transfected cells that are illuminated through the ultrasensitive luciferase reporter system. The present findings therefore demonstrate detectable brain-associated reporter activity rather than definitive brain-targeted delivery. Even with this distinction, the complete absence of detectable FLuc activity, compared with the measurable NLuc-GFP signal, illustrates how reporter sensitivity can alter conclusions about apparent expression in the brain and other tissues. The brain provided the most unique example; however, this effect was not limited to the central nervous system. NLuc-GFP also revealed reporter activity in the heart, kidneys, lungs, placenta (data not shown) and lymph nodes where FLuc signals were considerably weaker or near background (Fig 3D-E and 4D-E). These organs may be underappreciated when evaluating functional LNP distribution, particularly when relying primarily on FLuc imaging. In the heart, relatively low myocardial exposure to D-luciferin may restrict photon production when FLuc expression is modest. In the kidney, rapid renal clearance makes reporter output sensitive to the timing of substrate administration relative to image acquisition. Therefore, lower expression in the heart, kidneys, and lungs may also be underestimated when reporter abundance and substrate availability are insufficient to produce detectable FLuc output. This broader detection is important for both targeted and untargeted LNP development. The liver and spleen remained prominent with both reporters, consistent with their established roles in the uptake and clearance of systemically administered nanoparticles. However, the greater sensitivity of NLuc-GFP/FFz showed that reporter expression was not restricted to these dominant organs. This does not imply equivalent LNP delivery among tissues. Rather, it demonstrates that lower-expression organs can be classified as negative when the reporter–substrate system lacks sufficient sensitivity. A highly sensitive reporter may therefore reveal both desirable extrahepatic expression and off-target expression that would otherwise remain undetected.

The enhanced detection provided by NLuc-GFP reflects the integrated performance of the complete reporter system. Beyond its increased bioluminescence sensitivity, NLuc–GFP may function as a three-in-one reporter platform. NLuc/FFz enables sensitive whole-body and organ-level imaging, GFP permits fluorescence-based cellular and histological investigation, or GFP expression can be analyzed by flow cytometry to identify the specific cell populations undergoing functional mRNA delivery. Schaub et al. originally demonstrated the utility of the NLuc–GFP fusion for both sensitive in vivo imaging and fluorescence-activated cell analysis, supporting its application across organ and single-cell scales. Although cellular localization was not investigated in the present study, prominent lymph node-associated reporter signal was observed, warranting more detailed analysis of lymphatic expression. Future studies could use multiparameter flow cytometry of dissociated lymph nodes and other tissues to phenotype GFP-positive cells and determine which specific cell immune or stromal populations express the delivered mRNA. Complementary tissue-section analysis using direct GFP fluorescence or anti-GFP immunostaining could provide spatial confirmation of expression. These approaches would extend NLuc–GFP beyond organ-level biodistribution imaging to a multimodal platform capable of resolving both tissue-level and cell type-specific mRNA expression The present results also provide contrast for broader LNP-mediated protein expression reported in large-animal models. Ferraresso et al. detected translated protein across multiple organs following intravenous mRNA-LNP administration in swine, including the brain, heart, kidneys, and lungs (14). Together, these studies support the principle that apparent organ restriction can be influenced by reporter sensitivity. Differences in species, LNP composition, reporter design, dose, and detection methods prevent direct comparison; however, both studies indicate that functional expression may extend beyond the organs emphasized by conventional FLuc imaging.

Reporter stability represented a factor affecting longitudinal interpretation. The non-destabilized, NLuc-GFP reporter remained detectable for more than seven days, providing a broad imaging window and high sensitivity for low-abundance expression. However, persistent activity can obscure whether later luminescence reflects ongoing translation or previously accumulated protein. The addition of a degron motif, such as the PEST degron, reduced signal persistence, with NLuc-GFP-PEST returning to background by seven days. The stable reporter, long-acting luciferase is therefore better suited for maximizing detection and defining the broadest tissue profile, whereas the destabilized reporter provides improved temporal resolution with the same tissue clarity. Because the PEST-containing construct also produced lower signal at the earliest time point, its behavior may reflect differences in protein accumulation or steady-state reporter activity in addition to accelerated turnover. An additional contribution of this work was the development of a DSPE–PEG2000-based FFz formulation. Thin-film hydration produced an aqueous dispersion with high substrate recovery, which could be administered immediately after hydration. The formulation could also be lyophilized with trehalose and reconstituted before use (Supplemental Fig 2), potentially reducing day-of preparation and dependence on proprietary substrate formulations. Following reconstitution, the formulation retained a nanoscale size distribution with moderate changes in hydrodynamic diameter, polydispersity, and zeta potential. HPLC analysis provided an additional method for evaluating FFz recovery and chromatographic changes following lyophilization (Supplemental Fig 2). These findings establish the DSPE–PEG2000 formulation as an accessible approach for preparing FFz for in vivo imaging.

Overall, this study demonstrates that reporter sensitivity can substantially alter the interpretation of an mRNA-delivery experiment. Reporter mRNAs are central to preclinical LNP screening, but the apparent distribution of functional mRNA expression is ultimately constrained by the sensitivity of the reporter–substrate system. Across LNPs containing SM-102 or U105, both FLuc/D-luciferin and NLuc–GFP/fluorofurimazine identified the expected liver– and spleen-dominant expression pattern. However, NLuc–GFP produced substantially greater signal and detected low-level reporter activity in the kidney, heart, brain, and other tissues where FLuc activity was weak or undetectable. A weak or absent FLuc signal therefore should not be interpreted as definitive evidence of failed LNP delivery or mRNA translation. Instead, expression may remain below the detection threshold of the reporter system, particularly in tissues where substrate exposure is limited or transient. This work establishes two systems to provide a framework for evaluating both the detectable distribution and the duration of mRNA expression following LNP delivery with a cost-effective formulation of FFz.

## 4. Materials and Methods

### 4.1 Plasmid Design

The NLuc-GFP and NLuc-GFP-PEST plasmids were designed in SnapGene, purchased from GenScript (Nanjing, China), and inserted into the plasmid backbone, pGS-CMV-30A+30A+43A. The plasmid contains a polyA tract with two 30A segments and a 43A segment connected by small linkers to make a 103 polyA tail; see Supplemental Data for the complete annotated DNA sequences used.

### 4.2 Plasmid Transformation, Expansion, and Linearization

The plasmids obtained from GenScript were transformed into chemically competent NEB® Stable *E. coli* cells via heat shock (ice for 30 minutes, followed by 42 °C for 30 s), followed by recovery in NEB®-Stable Outgrowth Medium at 30 °C for 1 hour with shaking (250 rpm). Cells were plated on Luria-Bertani (LB) agar [Miller] containing kanamycin (50 µg/mL) and incubated overnight at 37 °C. Individual colonies were selected and expanded in 3–5 mL of LB broth containing kanamycin, then grown overnight at 37 °C with shaking (250 rpm). The cultures were then expanded into larger 1 L cultures using a 1:100 dilution and grown at 30 °C overnight. Plasmid DNA was isolated using the Qiagen EndoFree® Plasmid Maxi Kit (Qiagen, Hilden, Germany) according to the manufacturer’s instructions. Purified plasmid DNA concentration and purity were assessed by NanoDrop (ThermoFisher, Waltham, MA) with acceptable A260/A280 ratios >1.8. Plasmid integrity was further confirmed by agarose gel electrophoresis (0.9–1% agarose in TAE buffer); supercoiled plasmid DNA should be >90% quantified by gel densitometry. Plasmids were linearized downstream of the poly(A) tail using 12U / µg DNA BspQI-HF (New England Biolabs, MA, USA) in CutSmart buffer at 40 °C for 2 hours. Following digestion, linearized DNA was purified using spin column-based cleanup (NEB Monarch® High-Capacity DNA cleanup). Linearization was confirmed by agarose gel electrophoresis (0.9-1% agarose in TAE buffer). Whole Plasmid Sequencing was performed by Plasmidsaurus using Oxford Nanopore Technology with custom analysis and annotation to verify plasmids and polyA integrity.

### 4.3 In Vitro Transcription and mRNA Purification

mRNA was synthesized from the linearized plasmid DNA using the Takara IVTpro T7 mRNA Synthesis Kit (low dsRNA) (Takara Bio, Cat. No. 6134, Kusatsu, Shiga, Japan). Reagents were combined in the manufacturer’s recommended order, replacing uridine with N1-methylpseudouridine, co-transcriptionally capping with CleanCap® AG (3’OMe) (Trilink BioTechnologies, San Diego, CA), and the reactions were incubated at 37°C for 2 h. Following transcription, 4 µL of DNase I was added directly to each reaction, and the reactions were incubated at 37°C for 20 min. mRNA purified by lithium chloride precipitation, 30 µL nuclease-free water, and 30 µL LiCl precipitation solution were added to the DNase-treated reaction. Samples were mixed and incubated at −20 °C overnight, then centrifuged at maximum speed (> 20,000 g RCF) for 25 min at 4 °C. The supernatant was removed, and the RNA pellet was washed with 1.5 mL of chilled (−20 °C) 70% ethanol and centrifuged again at maximum speed for 15 min at 4 °C; this wash was repeated twice. After the final wash, pellets were briefly air-dried until opaque, then resuspended in nuclease-free water. Purified mRNA was stored at −80 °C until use.

### 4.4 mRNA Quantification and Characterization

Purified mRNA concentration and purity were measured using a Thermo Fisher NanoDrop spectrophotometer. RNA concentration was determined from absorbance at A260/A280. Preparations with an A260/A280 ratio near 2.0 were considered acceptable for downstream analysis. Transcript integrity and size distribution were evaluated by microfluidic capillary electrophoresis using an Agilent 2100 Bioanalyzer (Agilent Technologies, Santa Clara, USA) equipped with the RNA 6000 Nano Kit (Agilent Technologies, Cat. No. 5067-1511). Samples were diluted in RNase-free water to fall within the validated mRNA assay range of 25–250 ng/µL. Electropherograms and virtual gel images were evaluated for a predominant transcript peak or band at the expected size and for the absence of lower-molecular-weight fragments or degradation products. Only preparations exhibiting acceptable spectrophotometric purity and a predominant intact transcript were used for formulation and in vivo studies.

### 4.5 Lipid Nanoparticle Formulation, Encapsulation, and Characterization

Lipid nanoparticles were prepared using either SM-102 (BroadPharm, CA, USA) or U105 (Cayman Chemical, MI, USA) as the ionizable lipid. Both formulations contained ionizable lipid, cholesterol, DOPE, and DMG-PEG2000 at a molar ratio of 50:35:12.5:2.5. Lipids were dissolved in ethanol to prepare the organic phase. mRNA (Nluc-GFP, Nluc-GFP-PEST, or FLuc) was diluted in 10 mM citrate buffer (pH 4.0) to prepare the aqueous phase. The ionizable lipid-to-mRNA ratio was maintained at 10:1 by mass; the aqueous-to-organic ratio was kept at 3:1. The aqueous and organic phases were loaded into separate syringes and combined using a conventional syringe pump and a Herringbone Mixer Chip (Sigma-Aldrich, Cat. No. 926493), with a flow rate set to 10 mL/min. After formulation, LNP suspensions were transferred to 10-kDa molecular-weight-cutoff dialysis tubing and dialyzed overnight at 4 °C against 1 L of 1× phosphate-buffered saline (PBS), with one exchange after 4-6 h. Following dialysis, the LNPs were concentrated to the desired mRNA concentration/volume using 100-kDa molecular-weight-cutoff centrifugal filters. Hydrodynamic diameter, polydispersity index, and zeta potential were measured at room temperature using a Malvern Zetasizer Ultra Red (Malvern, PA, USA). Samples were diluted in 1× PBS for size measurements and in 10 mM HEPES for zeta potential measurements; all measurements were performed in triplicate. mRNA encapsulation efficiency was measured using the Quant-iT RiboGreen RNA assay (Thermo Fisher Scientific, Waltham, MA). Fluorescence was measured using a BioTek Synergy HT (Agilent, Santa Clara, CA) microplate reader set to (ex/em: 490/520), and encapsulation from intact LNPs was used to quantify unencapsulated mRNA. In contrast, total mRNA content was measured after disruption of the LNPs with 2% Triton X-100 in TE buffer.

### 4.6 Fluorofurimazine Substrate Formulation and HPLC Characterization

Fluorofurimazine (Ambeed, IL, USA) was incorporated into DSPE-PEG2000 (Ambeed) micelles by thin-film hydration. DSPE-PEG2000 and fluorofurimazine were combined at a 10:1 mass ratio. Specifically, 10 mg of DSPE-PEG2000 and 1 mg of fluorofurimazine were co-dissolved in 1-2 mL of chloroform in a 10 mL round-bottom flask. Chloroform was removed by rotary evaporation at 30 °C and 180 rpm. The resulting film was further dried under nitrogen until opaque and hydrated with sterile 5% dextrose at 50 °C with gentle agitation until a clear, homogeneous colloidal dispersion was obtained. The fluorofurimazine micelle formulation can be used directly following hydration without filtration, purification, or concentration. However, fluorofurimazine encapsulation was quantified by high-performance liquid chromatography using a ZORBAX SB-C18 column with a column guard (4.6 × 150 mm, 5 µm particle size). The mobile phase consisted of mobile phase A: 70% methanol (1% phosphoric acid) and mobile phase B: 30% water (1% phosphoric acid), delivered isocratically at 1 mL/min. The injection volume was 20 µL, and fluorofurimazine was detected at 430 nm. Under these conditions, fluorofurimazine eluted at approximately 1.801 min. A calibration curve was prepared using fluorofurimazine standards at 6.25, 12.5, 25, and 50 µg/mL. For analytical determination of micelle association and recovery, the formulation was processed through a 30-kDa molecular-weight-cutoff centrifugal filter. The micelle-containing retentate and free fluorofurimazine-containing filtrate were collected separately. The retentate was disrupted with methanol before HPLC analysis, then diluted to an appropriate concentration. Fluorofurimazine encapsulation and total recovery were both greater than 95%. For in vivo experiments, centrifugal filtration is not required and was only used for analytical characterization. For lyophilization, the DSPE–PEG2000/FFz thin film was hydrated at 50 °C with 10 mM HEPES containing 10% (w/v) trehalose, which served as the buffering agent and cryoprotectant, respectively. The hydrated formulation was transferred to amber glass vials and frozen at −80 °C for at least 6 h. Frozen samples were subsequently placed in a Labconco FreeZone Plus 2.5 L freeze dryer (Labconco, MO, USA) and lyophilized overnight to remove water. As a representative preparation, a formulation containing 1 mg FFz was initially hydrated with 1 mL of HEPES–trehalose buffer. Following lyophilization, the dried formulation was reconstituted with 0.5 mL of water preheated to 50 °C and gently rocked until a homogeneous dispersion was obtained. The formulation was characterized after lyophilization and reconstitution to evaluate changes resulting from freeze-drying. Hydrodynamic diameter and polydispersity index were measured by dynamic light scattering, and surface charge was assessed by zeta-potential analysis using the previously described conditions. Reconstituted samples were also analyzed by HPLC to quantify FFz without centrifugal filtration and to evaluate chemical degradation by monitoring the peak area and retention of the FFz peak, as well as the appearance of additional chromatographic peaks.

### 4.7 In Vivo mRNA Delivery and Bioluminescence Imaging

All animal procedures were performed in accordance with protocols approved by the Institutional Animal Care and Use Committee (IACUC). CD-1 mice (male/female) aged 6 to 12 weeks were used for all in vivo mRNA delivery and imaging studies. LNP-formulated mRNAs encoding codon-optimized Firefly luciferase (FLuc; TriLink Biotechnologies) or NLuc-GFP were prepared as described. Mice received 5 µg of LNP-formulated mRNA by retro-orbital injection. Bioluminescence imaging was performed using an IVIS Lumina XRMS imaging system (PerkinElmer, Hopkinton, MA, USA), 6 h post-injection. Mice were anesthetized with 3% isoflurane in oxygen during substrate administration, animal positioning, and image acquisition. Animals were positioned consistently within the imaging chamber. Field of view, binning, and f-stop were held constant across experimental groups, while exposure time was automatically determined to prevent detector saturation. Bioluminescent signal was quantified using Living Image software (PerkinElmer, Hopkinton, MA, USA).

For FLuc imaging, mice received 3 mg of D-luciferin potassium salt (GoldBio, Maplewood, MO, USA) via retro-orbital injection in a 30 mg/mL solution. For NLuc-GFP imaging, mice received a total dose of 0.20 mg of FFz, formulated in DSPE-PEG2000 micelles, via retro-orbital injection. Following live-animal imaging, mice were euthanized by carbon dioxide inhalation followed by cervical dislocation. The brain, heart, lungs, liver, spleen, and kidneys were collected immediately and arranged in a consistent orientation for comparing the two mRNAs for ex vivo imaging. Organ bioluminescence was measured using defined regions of interest and reported as total photon flux.

### 4.8 Longitudinal NLuc-GFP and NLuc-GFP-PEST Imaging

Reporter persistence was evaluated by comparing NLuc-GFP with NLuc-GFP-PEST. Mice received 3 µg of the respective LNP-formulated mRNA by retro-orbital injection and were imaged longitudinally at 6, 24, 48, and 168 h after mRNA administration using an IVIS imaging system. At each time point, mice received 0.20 mg FFz formulated in DSPE-PEG2000 micelles by retro-orbital injection. The same animals were imaged at each time point using consistent substrate doses, substrate-to-imaging intervals, animal positioning, field of view, binning, and f-stop. Exposure time was automatically selected. Whole-body bioluminescence was quantified by applying consistently sized regions of interest to each animal. The background signal was measured in an adjacent region lacking detectable bioluminescence. Total photon flux was used to compare reporter signals across constructs and imaging time points. Following live-animal imaging, mice were euthanized by carbon dioxide inhalation followed by cervical dislocation. The brain, heart, lungs, liver, spleen, and kidneys were collected immediately for ex vivo imaging. Organ bioluminescence was measured using defined regions of interest and reported as total photon flux.

### 4.9 Statistical analyses

The data were analyzed using descriptive statistics and presented as mean values ± standard deviation (SD) from independent measurements. The F-test was performed to confirm that the standard deviation between groups was not statistically significant before using a Mann-Whitney U-test to compare groups. The difference between variants was considered significant at p<0.05. All statistical analyses were performed using GraphPad Prism.

## Data Availability Statement

Data supporting the findings of this study are available from the corresponding author upon reasonable request.

## Acknowledgments

The authors express their gratitude to the OSU College of Pharmacy and to the Debra Larson O’Leary Research Fund in Pharmacy at Oregon State University for providing financial support. In addition, the authors would like to thank Namratha Turuvekere Vittala Murthy and the Sahay Lab for their support.

## Supplementary Materials

### This file includes

Supplementary Figure 1. DNA sequences for NLuc-GFP(-PEST) used in IVT

Supplementary Figure 2. DSPE-PEG2k/FFz Micelles Characterization Before and After Freeze-Drying

Supplementary Figure 3. Baseline bioluminescence following fluorofurimazine administration in the absence of reporter mRNA

Supplementary Figure 4. Whole-body and Ex vivo Biodistribution of SM102–NLuc–GFP–PEST mRNA LNPs

### Nluc-GFP DNA Sequence

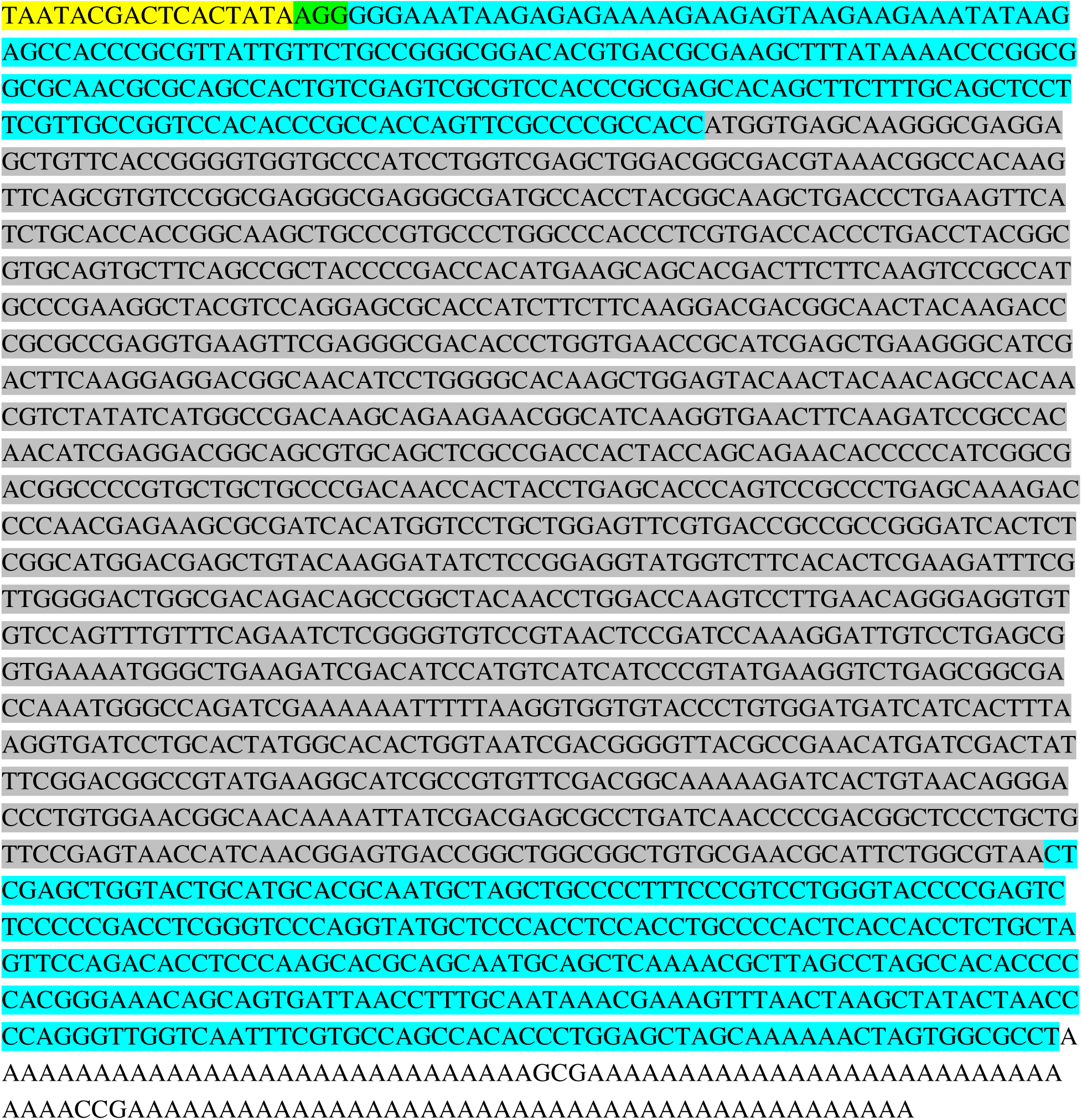

### Nluc-GFP-PEST DNA Sequence

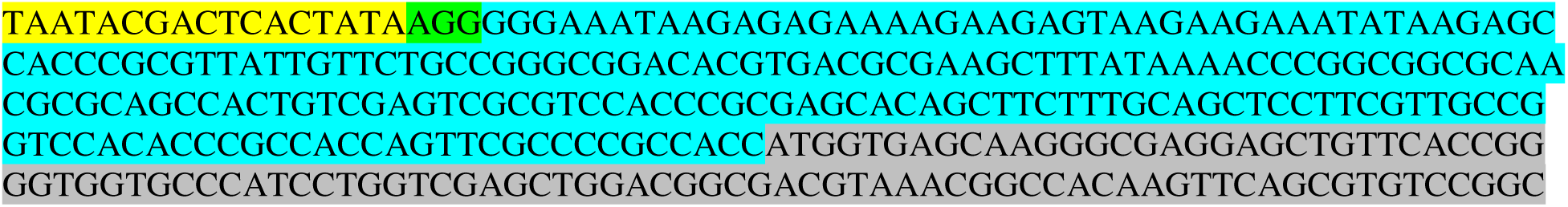
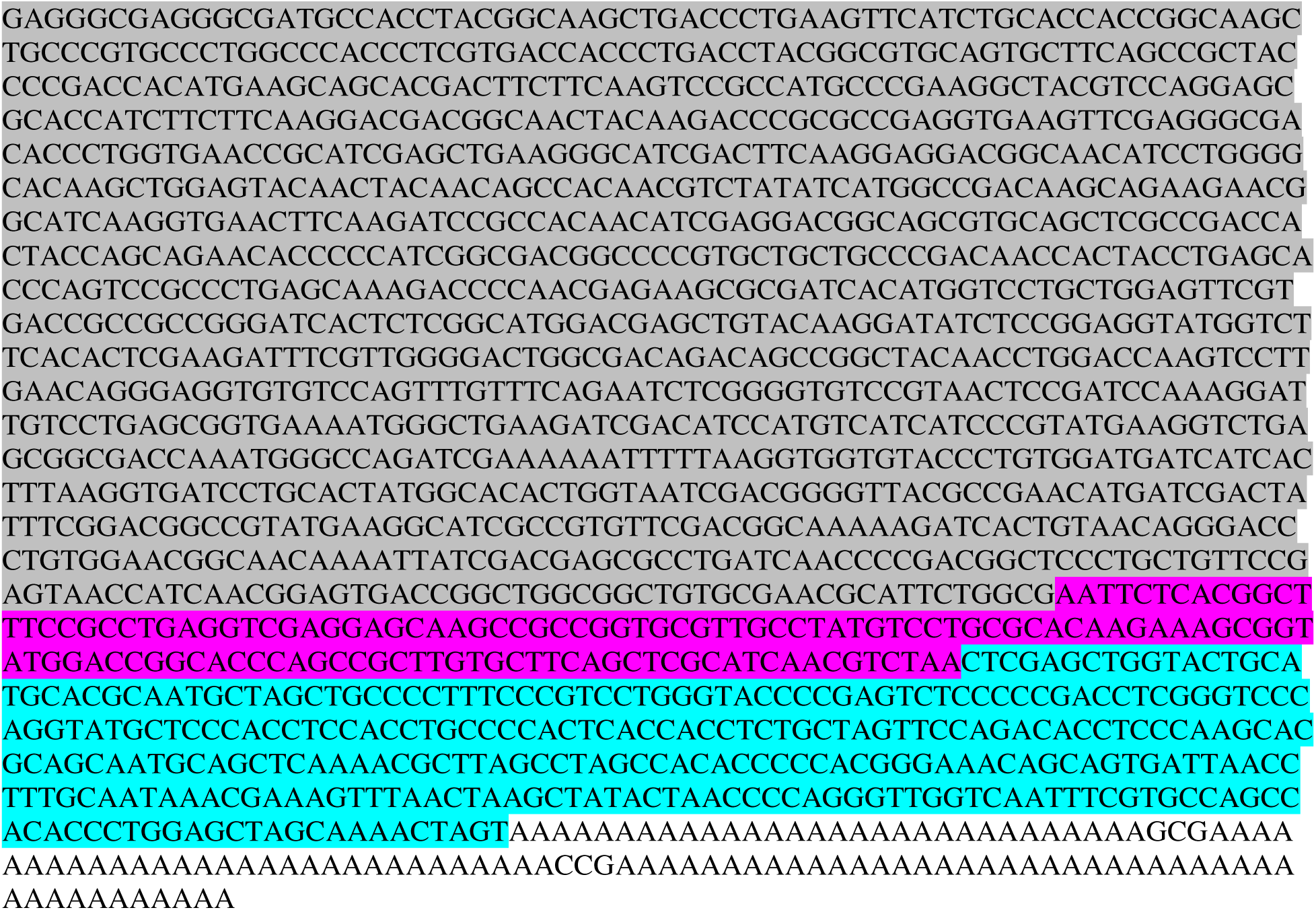

**Supplementary Figure 1.** DNA sequences of mRNA constructs. T7 promoter (yellow), CleanCap® Binding Sequence (Green), UTRs (Blue), CDS(gray), PEST signal (purple), PolyA(none)

**Supplementary Figure 2.**
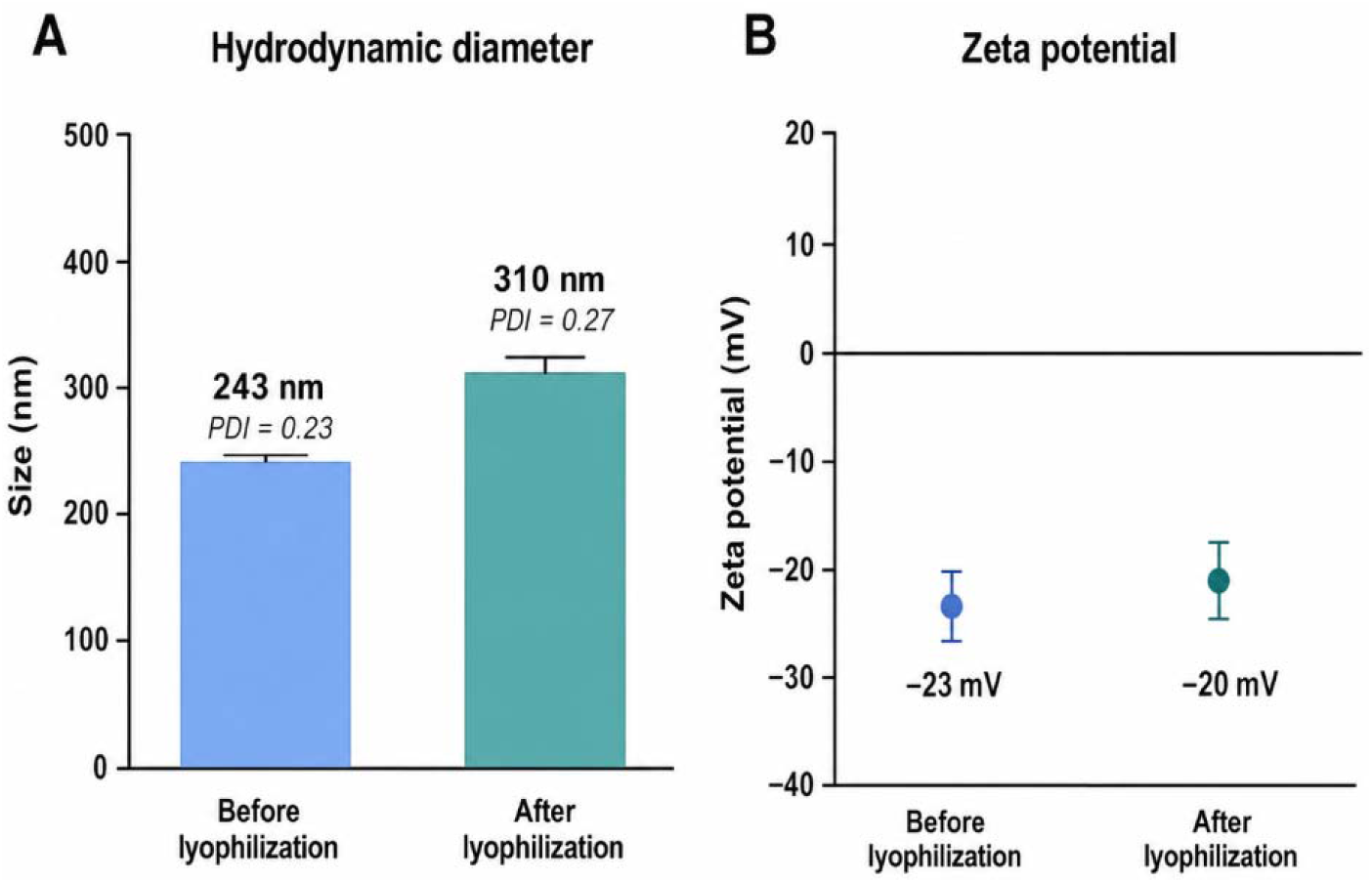
DSPE-PEG2k/FFz Micelles Characterization Before and After Freeze-Drying.

**Supplementary Figure 3.**
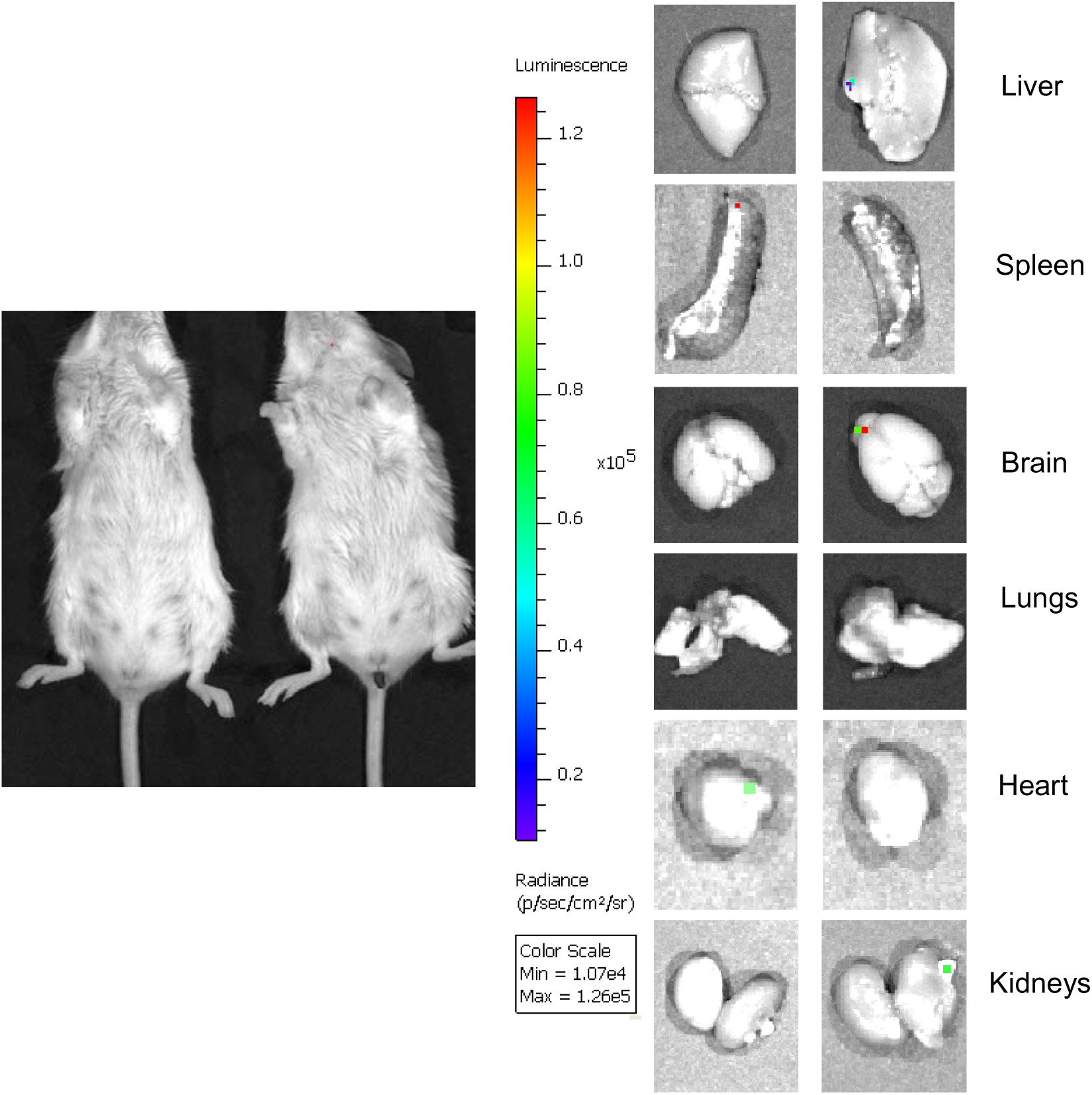
Baseline bioluminescence following fluorofurimazine administration in the absence of reporter mRNA. Control mice received fluorofurimazine (FFz) without NLuc-GFP encoding mRNA and were imaged by IVIS to determine whole-body background luminescence. Organs were subsequently collected and imaged ex vivo to assess tissue-associated background signal. (n=4)

**Supplementary Figure 4.**
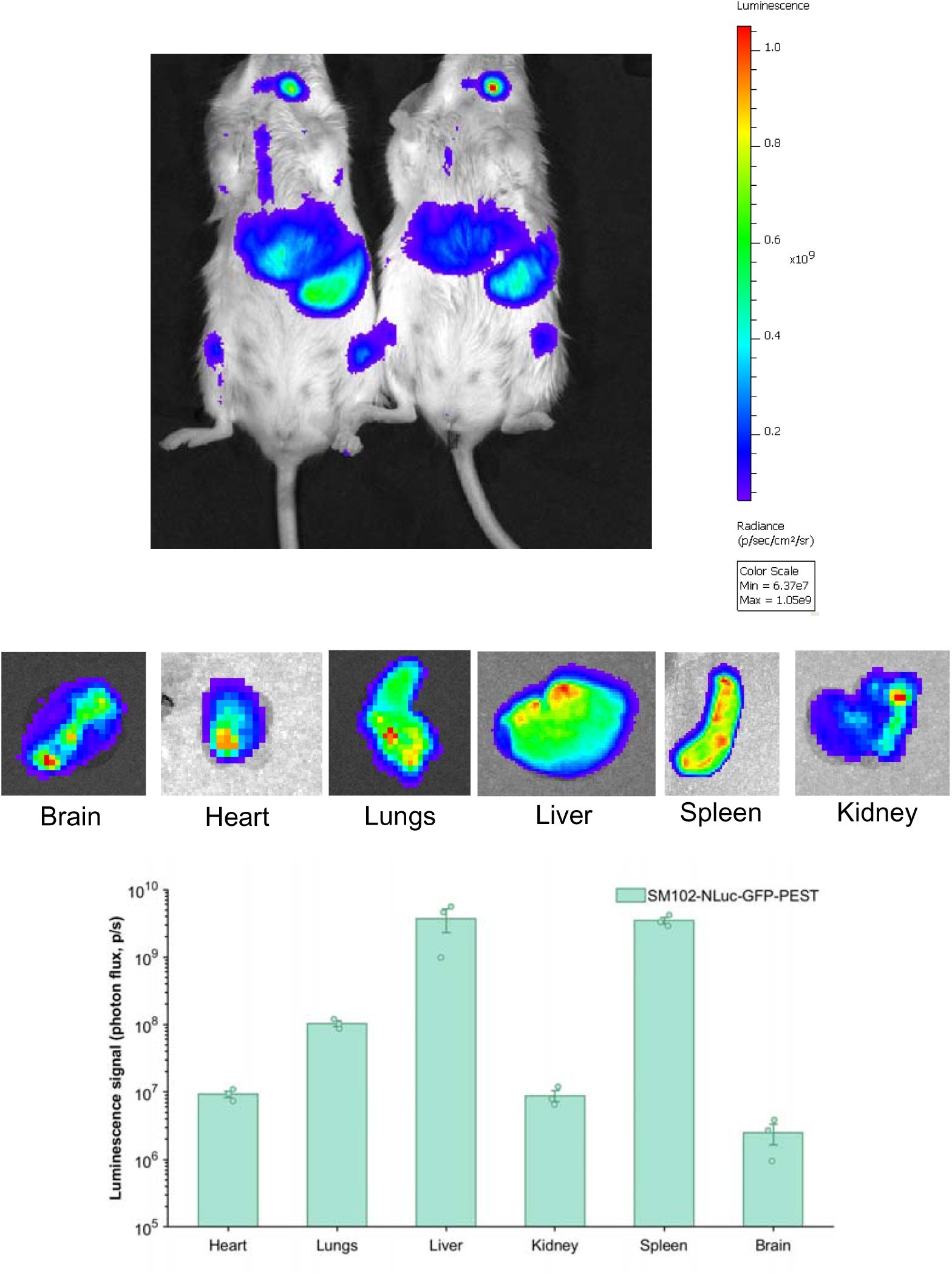
Whole-body and Ex vivo Biodistribution of SM102–NLuc–GFP–PEST mRNA LNPs. Mice received 3 ug SM102 LNPs containing NLuc–GFP–PEST mRNA and were imaged 6 h after administration following fluorofurimazine injection. Representative whole-body images and ex vivo images of the heart, lungs, liver, kidneys, spleen, and brain are shown. Organ luminescence was quantified as total photon flux. Bars represent mean ± SD, individual points represent separate animals (n=3).

